# Auditory Perception Induces Cortical and Thalamic Event-Related Desynchronization in the Mouse

**DOI:** 10.1101/2025.03.21.644604

**Authors:** Sarah H McGill, Qilong Xin, Taruna Yadav, Charlie W. Zhao, Patrick Paszkowski, Fabrizio Darby, Mrinmoyee Guha, Tramy Nguyen, David S. Jin, Yuval Nir, Jiayang Liu, Lim-Anna Sieu, Hal Blumenfeld

## Abstract

Studies of human perception have shown early cortical signals for primary information encoding, and later signals for higher order processing. An important late signal is the cortical event-related desynchronization (ERD) in the alpha (8-12Hz) and beta (12-30Hz) frequency band, which has been linked to human perceptual awareness. Detailed mechanistic investigation of the ERD would be greatly facilitated by availability of a suitable animal model. We conducted local field potential recordings in the mouse frontal association cortex (FrA), thalamic intralaminar centrolateral nucleus (Cl), primary auditory cortex (A1), and primary visual cortex (V1) during two auditory tasks. Fully audible brief 50 ms stimuli with both tasks produced early broadband gamma (30-100Hz) frequency activity at 0-250ms, followed by a late cortical alpha/beta ERD 250 – 750 ms after stimulus onset. The ERD was statistically significant in FrA and A1, but not in V1. Interestingly, a significant ERD was also observed in thalamic Cl. The magnitude of the ERD at full stimulus intensity, and the slope of the relationship between stimulus intensity versus ERD magnitude, were both largest in FrA, and smaller in Cl and A1. Conversely, for early broadband gamma activity the magnitude at full intensity and slopes were largest in A1, smaller in Cl and smaller still in FrA. These findings suggest that mice, like humans, process perceptual signals in hierarchically organized corticothalamic networks, and strongly support mice as a promising platform for further investigation of the ERD to better understand the origin and function of this robust yet understudied electrophysiological phenomenon.

**Significance Statement:** Auditory-induced alpha/beta event-related desynchronizations (ERDs), decreases in cortical activity between 8 and 30Hz following auditory stimulus presentation, are thought to represent systems underlying higher perception and cognition. However, fundamental mechanistic studies are difficult in humans, the dominant organism for studying this phenomenon. In this study we assess mice as a potential alternative model organism. Our results demonstrate that mice exhibit auditory-induced alpha/beta ERDs, that this response is also present in subcortical regions of the mouse brain, and that the cortical ERD is largest and most strongly related to auditory stimulus amplitude in association cortex rather than in primary auditory cortex. These results support the efficacy of mice as an ideal model organism for further examination of alpha/beta ERDs.

## Introduction

Conscious perception is a critical feature of normal sensory processing, and its disruption leads to behavioral and cognitive deficits (Blumenfeld, 2012; Giacino et al., 2014; Tononi & Edelman, 2000). Yet, the neurophysiological mechanisms of conscious perception remains elusive. Early electroencephalographic (EEG) and magnetoencephalographic (MEG) recordings in humans identified decreased cortical alpha (8-12Hz) and beta (12-30Hz) activity with both passive sensory stimuli and stimulus-response tasks (Pfurtscheller & Aranibar, 1977, 1979; for review see Pfurtscheller & Lopes da Silva, 1999). Subsequently, substantial evidence has related these alpha/beta band event-related desynchronization (ERD) responses with higher perception. In visual search and discrimination tasks, stimulus-induced ERD amplitude increases with number of distractors and with the similarity of target stimuli to distractors (Boiten et al., 1992; van Winsun et al., 1984). In memory tasks, lateral alpha/beta band ERDs are induced with memory retrieval, and haver higher amplitude in semantic versus episodic tasks (Klimesch et al., 1994), and in tasks of greater complexity (Dujardin et al., 1994). Bilateral transient alpha band ERDs are observed preceding and during un-cued volitional movements (Jin et al., 2024; Pfurtscheller & Aranibar, 1979). The hypothesis that attention partly underlies the ERD response (van Winsun et al., 1984) is supported by several studies (Bauer et al., 2014; Klimesch et al., 1997; Klimesch et al., 1998; MacLean & Arnell, 2011). Yet, data also support arousal mechanisms in the ERD, including auditory stimulation studies in sleep and anesthesia showing that early primary auditory cortex signals are relatively spared, whereas alpha/beta ERDs are markedly suppressed (Hayat et al., 2022; Krom et al., 2020).

Despite this important work, data identifying the specific networks responsible for alpha/beta ERDs are sparse. The central problem is that studies of alpha/beta ERDs, and of conscious perception more generally (Koch et al., 2016), are conducted mainly in humans, making mechanistic hypotheses testing difficult. Our goal was, therefore, to evaluate mice as a model for investigating sensory perception-related ERDs. In mice, techniques for measuring (Buzsáki et al., 2012; Cardin et al., 2020; Huang et al., 1996; Steinmetz et al., 2018) and manipulating (Cardin, 2012; Grossman et al., 2017; Jackson et al., 2016; Lim et al., 2013) activity of specific neuronal populations are well established. Validity of the mouse model is strengthened by mouse behavioral indicators of stimulus perception (Carandini & Churchland, 2013; Kolata et al., 2007) and by data showing early components of sensory processing similar to humans (Henry, 1979; Laramée & Boire, 2015; May, 2008; Moore & Moore, 1971; Vézina, 2013). However, to our knowledge no previous studies have assessed whether mice exhibit alpha/beta ERDs with sensory perception.

Towards this end, we conducted local field potential (LFP) recordings of mouse frontal association cortex (FrA), centrolateral thalamic nucleus (Cl), primary auditory cortex (A1), and primary visual cortex (V1) during two tasks with variable auditory stimulus intensities. This approach allowed us to characterize responses in mouse analogs of human cortical and thalamic regions involved in primary or higher order sensory perceptual processing. We found that mice exhibit delayed alpha/beta ERDs in FrA, Cl, and A1 between 250 and 750 ms following auditory stimulus presentation. The ERD was stronger and more directly related to stimulus intensity within FrA than in any other region measured. Moreover, the FrA, Cl, and A1 exhibited rapid, broadband increases in LFP power between 0 and 250 ms following auditory stimulus presentation, strongest in A1. Finally, we confirm that the mouse visual cortex shows a weak but statistically significant cross-modal early gamma response to auditory stimulus presentation without the alpha/beta ERD. Ultimately, these findings broadly highlight the potential for mice as an effective platform for examining the mechanisms of conscious perception, and specifically provide a promising animal model for further investigation of the fundamental nature of the ERD.

## Methods

### Animal Research Ethics

To ensure that animals used in this study received humane treatment, all protocols administered to animals were reviewed and approved by the Yale Institutional Animal Care and Use Committee (IACUC) before implementation. In addition, annual inspections were performed by the Yale IACUC to independently verify ethical standards were met.

### Mice and Surgical Procedures

All mice employed in this study were C57Bl/6 mice from Charles River. At the time of surgical manipulation, the animals were between 2 and 6 months of age. In total 32 mice were employed in this study, of which 16 were male and 16 were female.

Mice were anaesthetized for surgery using isoflurane inhalation (3% saturation; Covetrus) followed by intraperitoneal injection with a ketamine (90 mg/kg; Covetrus) and xylazine (9 mg/kg; Covetrus) mixture. As ketamine reduces core temperatures in rodents (Wixson et al., 1987), the mice were warmed to 37°C using a heating pad and heat therapy pump (Gaymar T/Pump Localized Therapy System). To provide analgesia both during and after surgical operations, mice were injected subcutaneously with Buprenorphine XR (3.25 mg/kg; Ethiqa) before surgery.

To record neural signals from target regions, mice were implanted with 0.15mm outer diameter insulated stainless steel bipolar twisted-pair local field potential electrodes (E363/3-TW/SPC; Protech International) as well as 1.60mm shaft diameter ground screws (8IE3639616XE; Protech International). To permit head fixation, the mice were additionally implanted with a custom-fabricated titanium headplate. Stereotaxic coordinates of electrode locations were calculated relative to bregma using The Mouse Brain in Stereotaxic Coordinates Second Edition (Franklin & Paxinos, 1997). For each mouse the electrodes were targeted to up to three of either the frontal association cortex (+2.58mm AP; +/− 1.39mm ML; −1.40 mm SI), the centrolateral intralaminar thalamic nucleus (−1.58mm AP; +/− 0.65mm ML; −2.72 mm SI), the primary auditory cortex (−2.92mm AP; +/− 3.83mm ML; −2.22mm SI), or the primary visual cortex (−3.40mm AP; +/− 2.47mm ML; −1.11mm SI). A ground screw was targeted to anterior parietal lobe pia, and the headplate was adhered to the skull above the lambdoid sutures.

To insert these implants, mice were shaved on their scalps and then mounted onto a stereotaxic frame (Kopf Model 962 Dual Ultra Precise Small Animal Stereotaxic Instrument). Their scalps were then cleaned using alcohol wipes, disinfected with betadine, and injected subcutaneously with lidocaine (0.10 mL at 2% wt/wt; Covetrus) before being excised between the posterior section of the frontal bone plates and anterior section of the interparietal bone plates. The surface of the skull was leveled until the SI positions of bregma, and lambda were within 0.10mm of each other after which the final locations of both craniometric points were recorded. The target positions of the electrodes and ground screws along AP and ML axes were located using the stereotaxic arm and marked on the skull surface. Burr holes were then made at these locations using a handheld micromotor electrical drill (Rampower). With the brain exposed, the electrodes were inserted to their target depths using a stereotaxic arm as a guide, and the ground screw was driven into its burr hole using a screwdriver to a depth of 1mm. Dental cement (C&B) was applied to both the electrodes and the ground screws to ensure they were secure.

The headplate was applied to the posterior section of the exposed skull, leveled with a bubble level, and cemented in place. Cement was applied to the edge of the exposed scalp to create a seal between the scalp and skull and prevent infection. Exposed wires from the electrode and ground screws were inserted into threaded cylindrical connectors (Protech International) for interfacing with electrophysiology equipment and covered with acrylic resin for strength and protection (Jet Denture). Postoperatively, mice were injected with carprofen (brand name Rimadyl; 5 mg/ml; Zoetis) daily for up to three days to prevent inflammation.

### Auditory Stimuli and Behavioral Tasks

Mice were trained in two auditory perception tasks, which both included a brief 50 ms auditory stimulus presented over a range of sound intensities superimposed over a white noise background. Sounds were presented to the mice using speakers (HiVi Tn25 Fabric Dome Tweeter) enclosed in 3D-printed housings flanking the running wheel. Stimuli were generated using a dedicated digital to analog converter and amplifier circuit board (Geekworm X400 V3.0 Audio Expansion Board). The primary auditory stimulus was a complex chord consisting of 17 constant, falling and/or rising sine wave tones distributed between 4KHz and 20KHz frequencies overlaid on each other, with a duration of 50ms. These stimulus features were chosen to ensure the auditory cortex, which in mice is tonotopically organized (Stiebler et al., 1997), undergoes spatially widespread activation with above-threshold stimulus presentation. White noise was played continuously as a background mask and to minimize the effects of natural variation in ambient noise. Intensities of the complex chord stimuli and white noise background were measured using a calibrated class 1 sound level meter (DSM403SD; General). The median volume of the ambient noise in the behavioral testing area was 41.6 dB. The median volume of the audio system playing the white noise mask alone was 63.5 dB. The median volume of the complex chord stimulus played at maximum intensity with the white noise mask was 65.7 dB. For both behavioral tasks the complex chord stimulus was played over a range of fixed intensities modulated by an amplitude scalar, a unitless multiplier applied to the amplitudes of audio file waveform samples to control playback volumes. This scalar ranged from a value of 0 (no stimulus, corresponding to background white noise being played alone at 63.5 dB), to 0.1 (maximum intensity stimuli, resulting in a total measured volume of 65.7 dB including background white noise).

Behavioral paradigms were administered in automated fashion using a dedicated single board computer (Raspberry Pi 4 4GB) running custom python software. Using the built-in 5V GPIO terminals, this computer was wired directly to the CED 1401 data acquisition unit to allow the exact timing of all events associated with the behavioral paradigms to be recorded. The two behavioral tasks were a passive listening task where sound stimuli were presented and no response was required, and an auditory go/no-go task where mice were trained to respond to sound stimuli by licking a lick port to receive a sucrose water reward (5% sucrose). Initial analysis of the electrophysiological data from the two tasks showed very similar early broadband activity, and late alpha/beta ERDs for both tasks, so data from the two tasks were combined in the final analyses to maximize sample sizes. Individual mice either underwent the passive listening task alone, the auditory go/no-go task alone, or both tasks in sequence.

Mice were allowed a minimum of 3 days to recover from surgery before acclimatization to the testing apparatus with head fixation while on a running wheel for 30 minutes daily over a minimum of 2 additional consecutive days. The passive listening task required no additional training, but the auditory go/no-go task required an average of 4-5 weeks of training for mice to associate the sound stimulus to the sucrose water reward. Passive listening task sessions ended once 45 minutes had passed, while go/no-go task sessions ended once 45 minutes had passed or the mice had received its daily allotted quota of water. In both tasks, individual trials consisted of a 2-8s intertrial interval, a 50ms sound stimulus phase, and a fixed 1.5s post-stimulus delay phase. The 2-8s intertrial intervals were characterized by a delay of random length distributed along an exponential curve with a median duration of 4s. The stimulus phases were characterized by presentation of either no auditory stimulus, or presentation of the 50 ms complex chord stimulus at a random volume within the range described above. In the passive listening task the next trial began immediately after the 1.5s post-stimulus delay, whereas in the go/no-go task there was an additional interval of 3.5 s for licking during successful trials and 13.5 s for punishment in unsuccessful trials, before the next trial would commence. Electrophysiological data for both tasks were initially cut into epochs spanning from 1.5s before to 1.5s after each stimulus, and final analyses were performed only on data from 1s before to 1s after each stimulus, so that identical peri-stimulus intervals were analyzed for both tasks.

### Electrophysiology

Local field potential waveforms were amplified by 1000x (A-M Systems Model 1800 Microelectrode AC Amplifier; 0.1Hz low cutoff, 10kHz high cutoff), before being low pass filtered at 100Hz using an analog filter (Model 3364 Krohn-Hite). To minimize the effects of mechanical or electrical interference on these recordings, all electrophysiological measurements were made with mice inside a Faraday cage grounded to the building and mounted to an air table. The electrophysiological waveforms were recorded using a CED data acquisition unit (Micro1401-4 with ADC12 expansion unit; CED) at a sampling rate of 1000Hz.

### Histology

Histology was employed to verify the locations of the electrodes. For this purpose, mice were anaesthetized with isoflurane inhalation and intraperitoneal injection of a ketamine and xylazine mix. They were then euthanized through intraperitoneal injection of 0.05mL Euthasol (Virbac). To fix the brain tissue, the mice were administered intracardial perfusions using a solution of heparin (1000 units/ml; Sagent Pharmaceuticals) in phosphate buffered saline (PBS; Sigma-Aldrich), followed by a mixture of formaldehyde (4% wt/wt; Thermoscientific) in PBS. The brains were then excised and allowed to sit in a formaldehyde solution for a minimum of 24 hours. After fixation, they were washed in PBS and coronally sectioned into 60µm slices using a vibratome (Leica vt1000s). The slices were mounted onto polarized slides (Epredia) and allowed to dry for 24 hours before being stained with cresyl violet (FC Neurotechnologies). A glass coverslip (Epredia) was secured on top of the stained slices with Cytoseal 60 (Epredia) for protection. The slides were imaged using a Leica DM6 B microscope. All electrophysiological recordings from electrodes implanted in an incorrect location were excluded from analyses.

### Data Preprocessing

All data were analyzed using MATLAB (R2023b; Mathworks). The MATLAB/SON interface library provided by CED was used to open and read from the electrophysiological recording files in MATLAB. Individual trials with LFP recording amplitudes over 500µV in at least one sample were rejected as outliers. For any condition, only animals with at least ten trials applicable to the condition were included in analyses to maximize the signal-to-noise ratio in the results. Exact timings of auditory stimuli were determined by analyzing the audio channel waveforms generated by the digital to analog converter and amplifier board and recorded by the CED data acquisition unit. For this purpose, waveform elements exceeding 3.5 standard deviations from the median amplitude and occurring within 10ms of each other were clustered. Clusters shorter than 0.03s or longer than 0.07ms, or those occurring outside the 1.55s span following a TTL pulse marking execution of the stimulus presentation function, were considered erroneous and excluded. The stimulus start time was defined as the timestamp of the first sample in each valid cluster. In trials where no stimulus was presented or when low-intensity stimuli were indistinguishable from background noise, a fallback estimate of 119.3ms after stimulus function execution was used. This estimate was derived from diagnostic tests measuring the median delay between the TTL pulse and stimulus onset for maximum-intensity complex chord stimuli. After preprocessing, recordings from each trial were cut into 3-second epochs spanning the 1.5 seconds before and after the auditory stimulus onset.

### Time-Frequency Analysis

Time-frequency spectrograms were computed from LFP recordings aligned to stimulus onset. Using a fast-Fourier implementation (*spectrogram* function in MATLAB) of a discrete Fourier transformation (DFT), power was computed in 250ms time windows with 218ms overlap. The spectrogram power values at each time point and frequency were converted to decibel scale (10×log_10_(Power/Baseline)), where baseline was defined for each frequency as the mean power over the interval from 500ms prior to auditory stimulus onset until the point just prior to stimulus onset. These time-frequency spectrogram values were then pooled within each animal by averaging across trials at each stimulus intensity for downstream analysis.

To identify statistically significant time-frequency changes associated with presentation of auditory stimuli, we compared maximum-intensity trials (65.7 dB total, with 50 ms maximum intensity complex chord stimulus plus white noise background mask) to no-stimulus trials (63.5 dB, with white noise background mask only). Statistical significance of the differences between these two conditions were determined using a cluster-based permutation test (see next section). The time-frequency response associated with stimulus presentation in each recorded region was identified by calculating the difference between the maximum-intensity and no-stimulus conditions taken as the mean across mice (Figure 1).

**Figure 1.**
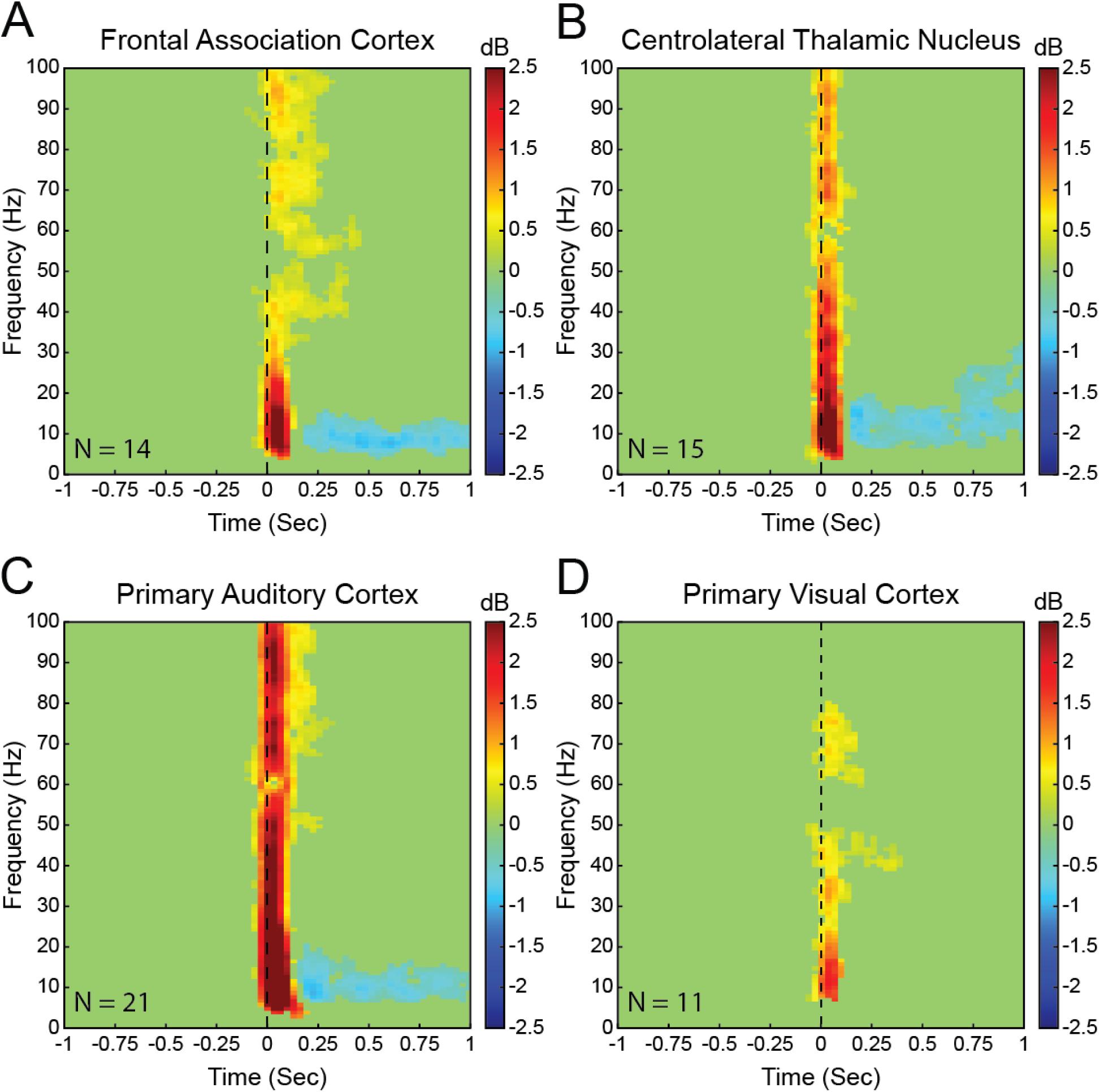
Time-frequency analysis of local field potential recordings in target brain areas following presentation of auditory stimuli at the maximum intensity (65.7 dB) tested minus no stimulus. Peristimulus spectrograms were identified for all maximum intensity and no-stimulus trials, decibel normalized to the prestimulus baseline (the 500ms before stimulus presentation). The mean spectrogram time-locked to stimulus onset (time 0, vertical dashed line) was calculated for each mouse and target location pair. Statistically significant differences (p<0.05, 5000 iterations of cluster-based permutation analysis) between the mean maximum intensity auditory stimulus trial spectrograms and the mean no stimulus trial spectrograms across mice were calculated and are displayed here. Frontal association cortex (A), centrolateral thalamic nucleus (B), and primary auditory cortex (C) show early increases in alpha (8-12Hz), beta (12-30Hz), and gamma (30-100Hz) band frequencies with late decreases in alpha and beta frequency bands. Primary visual cortex (D) shows only increases to early alpha, beta, and gamma band power. Number of mice (N) are shown for each panel.

### Cluster-Based Permutation Testing

Maximum stimulus-intensity peristimulus spectrograms were compared to no stimulus trial spectrograms using hypothesis-free cluster-based permutation testing. This approach has been described previously (Groppe et al., 2011; Kronemer et al., 2022). Briefly, a null distribution was created by randomly shuffling data between the maximum stimulus intensity and the no stimulus conditions across mice (n=number of animals). A first level two-tailed t-test was performed comparing maximum stimulus intensity and no stimulus conditions, and the associated t-values were calculated for each time point and frequency. These t-values were used to cluster the data based on frequency and temporal adjacency, defined as the neighboring frequency and time points respectively. Only those clusters whose sum of t-values were highest were kept for each iteration. This shuffle and clustering process was repeated with 5000 iterations to create a null distribution. Finally, a two-tailed t-test was performed on the actual data of the maximum-intensity and no-stimulus intensity conditions and similarly clustered based on frequency-time adjacency. Only those clusters with sum of t-values higher than the 95^th^ percentile of the null distribution were kept as statistically significant. This approach was designed to address the multiple comparison problem associated with comparing data across large numbers of time points and frequencies. A modified version of the clust-perm1 function from the MATLAB Mass Univariate ERP Toolbox created by David Groppe (Groppe et al., 2011) was employed to conduct this analyses. Finally, to remove small spurious clusters, only those clusters exceeding 89 time-frequency elements (1% of the total number of time-frequency elements in the spectrogram) in size were kept.

### Power Spectral Density Analysis

Power spectral densities (PSDs) of peristimulus recordings were obtained for each trial using Welch’s method applied to two time-windows of interest: the early post-stimulus period (0-250ms following stimulus onset) and the late post-stimulus period (250-750ms post-onset). This same method was also used to calculate the PSDs of the prestimulus baselines (defined as the 500ms prior to stimulus onset). These computations were performed using a 100 ms Hamming window with 50% overlap between segments. The PSDs were next aggregated within each animal by calculating the mean PSD for each target brain location, time window, and sound intensity across all relevant trials. The PSDs of each type for each animal were then decibel normalized (10×log_10_(Power/Baseline)) to their corresponding mean baseline PSDs, with the baseline interval as already defined. The no-stimulus condition PSDs were then subtracted from their corresponding maximum-intensity stimulus condition PSDs for each combination of brain location and time window. Finally, the PSDs were combined across animals by identifying the average PSD difference between maximum intensity and no-stimulus conditions within each brain location and time window (Figure 2).

**Figure 2.**
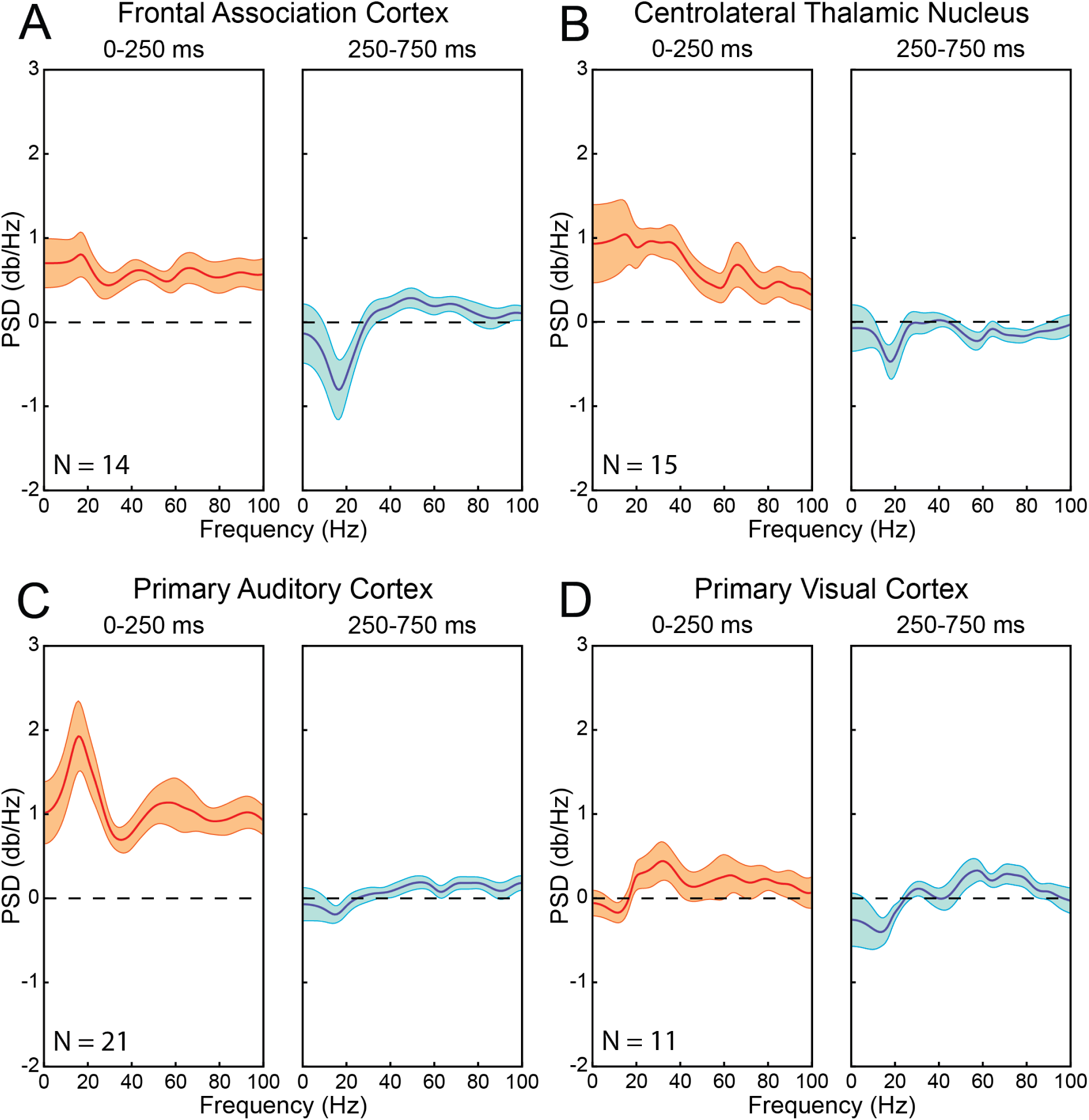
Power spectral density (PSD) curves of local field potential (LFP) recordings in target brain regions during early (0-250ms after stimulus onset; left graphs of each panel) and late (250-750ms after stimulus onset; right graphs of each panel) post-stimulus response periods of trials with auditory stimuli presented at the maximum intensity (65.7 dB) minus no-stimulus trials. PSD curves were calculated for early and late post-stimulus response periods in all maximum intensity trials. The PSDs were decibel normalized to their respective prestimulus baselines. The average PSD across all trials was identified for each mouse and target region pair, and then the mean PSDs across all mice were calculated and are displayed here. Presentation of auditory stimuli results in broadband increases to mouse frontal association cortex (A), centrolateral thalamic nucleus (B), and primary auditory cortex (C) with peaks in alpha (8-12Hz) and beta (12-30Hz) frequency bands during the early post-stimulus period. In the late post-stimulus period, prominent alpha-beta desynchronizations are found in the frontal association cortex and centrolateral thalamic nucleus, with smaller changes in primary auditory cortex. Primary visual cortex (D) shows limited increases to LFP power in mainly beta and low gamma (30-100Hz) bands during the early post-stimulus period. Center lines plot mean of PSDs across mice. Upper and lower bands show standard error of the mean. Number of mice (N) are shown for each panel; same animals and trials as in Figure 1.

### Stimulus Intensity Regression

To identify the effect of stimulus intensity on the spectral composition of electrophysiological responses elicited from stimulus presentation, a regression analysis of selected time-frequency bands against stimulus intensity was performed. The selected time-frequency bands consisted of the gamma band (30-100Hz) of the early post-stimulus period (0-250ms) and the alpha/beta band (8-30Hz collectively) of the late post-stimulus period (250 to 750ms). Towards this end, the subset of the peristimulus time-frequency spectrograms within the selected time-frequency bands were extracted for each trial following the baseline-corrected decibel normalization as detailed above. We then calculated the mean values across all frequency and time points within the selected time-frequency bands for each trial. Following this, the mean early gamma band and the mean late alpha/beta band responses were calculated within each animal for each target brain region across all trials of the same intensity. Linear models were fit to stimulus intensity versus either early gamma band power or late alpha/beta band power for each target brain region, treating mice as replicates (see Fig 3). The strength of this relationship was identified by calculating Spearman’s rank correlation coefficient, and the slope of the linear fit. The significance of fit was identified by calculating the p-value associated with the coefficient of linear regression.

**Figure 3.**
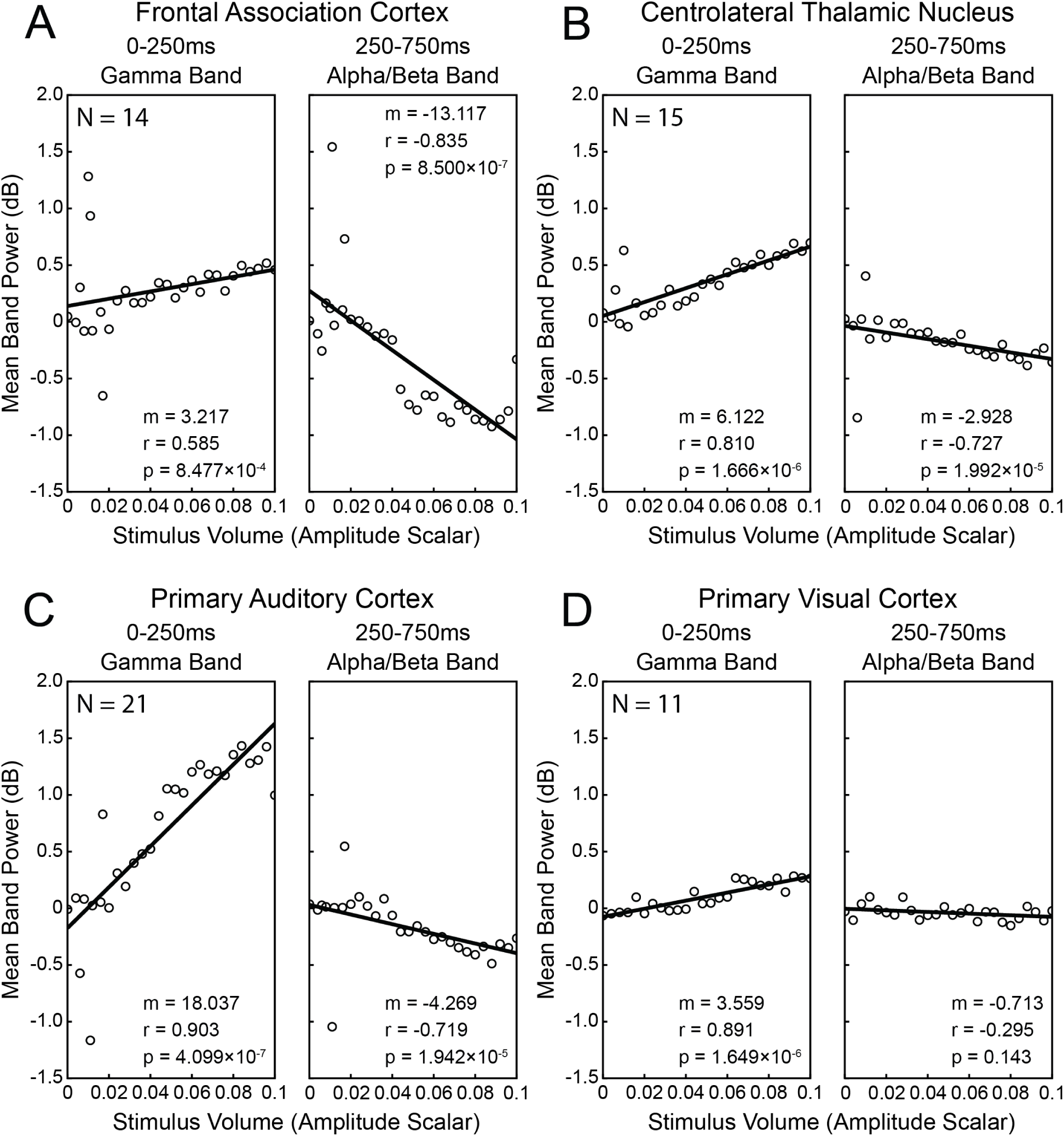
Regression of LFP power against stimulus intensity for early (0-250ms) post-stimulus gamma (30-100Hz) frequency band and late (250-750ms) alpha/beta (8-12Hz and 12-30Hz) frequency bands in all target brain regions. For each trial, the band power of the early post-stimulus gamma band and late post-stimulus alpha/beta bands were decibel normalized against the prestimulus baseline (the 500ms prior to stimulus onset). The average band power changes versus baseline at each stimulus intensity were then calculated within each mouse, and the mean of these values across mice is displayed here. Stimulus intensity values shown here are scalar multipliers of the 50ms complex chord audio waveform samples, and are in arbitrary units such that 0 corresponds to no sound, and 0.1 corresponds to the maximum sound stimulus used. Note that a white noise background mask was also played at all times. Mouse frontal association cortex (A), centrolateral thalamic nucleus (B), primary auditory cortex (C), and primary visual cortex (D) exhibit significant positive relationships between early post-stimulus gamma power and stimulus intensity as well as between late post-stimulus alpha/beta power and stimulus intensity, although the strength of these relationships varies. Spearman correlation coefficient (r). Significance of correlation (p). Regression slope (m). Number of mice (N) are shown for each panel; same animals as Figures 1 and 2 but using variable stimulus intensity trials.

## Results

To characterize mouse brain activity across cortical and subcortical regions during auditory perception, we measured local field potentials (LFPs) within the frontal association cortices (FrA), centrolateral thalamic nuclei (Cl), primary auditory cortices (A1), and primary visual cortices (V1) of mice during passive listening and auditory go/no-go tasks featuring stimuli of variable intensity. We then compared across mice the mean spectral composition of the electrophysiological recordings made during trials with maximum-intensity (65.7 dB with both stimulus and white noise background) stimuli to those made during no-stimulus (63.5 dB with white noise background only) control trials pooled between both tasks. This approach revealed that presentation of auditory stimuli results in significant (p<0.05; permutation test) changes to LFP power in all brain regions targeted (Figure 1).

FrA, Cl, and A1 showed temporally bimodal responses with distinct early (0-250ms) and late (250-750ms) response components. The early response components were characterized by increases in LFP power spanning alpha (8-12Hz), beta (12-30Hz), and gamma (30-100Hz) frequency bands. Following stimulus onset and up to 250ms post-stimulus, the FrA (Figure 1A) exhibited a +0.98 dB mean increase in alpha band power, a +0.69 dB increase in beta band power, and a +0.44 increase in gamma band power. The Cl (Figure 1B) in this same time frame showed a +0.78 dB increase in alpha band power, a +0.76 dB increase in beta band power, and a +0.53 dB increase in gamma band power. And the A1 (Figure 1C) in the early response period featured a +1.84 dB increase in alpha band power, a +1.49 dB increase in beta band power, and a +1.08 dB increase in gamma band power. The late response components within these regions consisted of a decrease to both alpha and beta band power that is characteristic of alpha/beta event-related desynchronization (ERD). The FrA (Figure 1A) during this period showed a −0.80 dB decrease to alpha band power and a −0.24 dB decrease to beta band power. The Cl (Figure 1B) in this same interval exhibited a −0.52 dB decrease to alpha band power and a −0.35 dB decrease to beta band power. And the A1 (Figure 1C) during the late response period featured a −0.59 dB decrease to alpha band power and a −0.24 dB decrease to beta band power.

Interestingly, the V1 (Figure 1D) also exhibited a response to presentation of maximum-intensity auditory stimuli significantly different from no-stimulus controls. Like the FrA, the Cl, and the A1, the V1 featured a broadband increase in LFP band power within 250ms of stimulus onset. Yet, the increase in LFP band power observed in the V1 was weaker than those found in other regions. During the early post-stimulus response period V1 showed a +0.06 dB increase in alpha band LFP power, a +0.18 dB increase in beta band LFP power, and a +0.28 dB increase in gamma band power. Moreover, no significant difference in LFP band power was identified between maximum-intensity stimulus trials and control no-stimulus trials following this initial response.

To visualize the frequency composition of changes in LFP within the target brain regions and time windows of interest following stimulus presentation, we calculated the mean power spectral density (PSD) across mice of the electrophysiological recordings made during maximum-intensity stimulus trials minus no stimulus trials for the early (0-250ms) and late (250-750ms) post-stimulus periods. In the early post-stimulus period the FrA (Figure 2A, left), Cl (Figure 2B, left), and A1 (Figure 2C, left) showed broadband increases to LFP PSD compared to intertrial interval baselines. The mean change in LFP PSD across all frequencies in the FrA, Cl, and A1 were found to be +0.60 dB/Hz, +0.86 dB/Hz, and +1.22 dB/Hz respectively. In addition to the broadband changes, these regions showed peaks in the alpha and beta frequency ranges. The greatest magnitude of the early LFP post-stimulus response in the FrA was found to be at 17 Hz with a PSD peak of +0.80 dB/Hz, in the Cl was identified as 15Hz with a peak at +1.04 dB/Hz, and in the A1 was determined to be 16 Hz with a peak at +1.93 dB/Hz. In V1 (Figure 2D, left), there were also increases in LFP PSD during the early post-stimulus response period, although these increases were limited mainly to the beta and low gamma frequency bands, with a peak of +0.44 dB/Hz at 32Hz.

In the late post-stimulus period, FrA (Figure 2A, right) and Cl (Figure 2B, right) showed substantial decreases to alpha and beta band LFP PSD. Within this time span, the response was not broadband, but instead localized to lower frequencies. Prominent minima were found during the late post-stimulus period in the FrA at 17 Hz with a decrease of −0.81 dB/Hz and in the Cl at 18 Hz with a decrease of −0.47 dB/Hz, and a smaller minimum was found for A1 (Figure 2C, right) at 15 Hz with a decrease of −0.19 dB. In V1 (Figure 2D, right), we observed a minimum at 14 Hz with a decrease of −0.40 dB/Hz in the late post-stimulus period, although this change did not reach statistical significance with time-frequency permutation-based clustering analysis.

We presented stimuli of varying intensities to mice during both passive listening and go/no-go tasks. To determine the effect of stimulus intensity on the magnitude of the responses observed in the target brain regions, we calculated for each trial the mean gamma band power of the peristimulus LFP recordings during the early post-stimulus period, as well as the mean alpha/beta band power of the recordings during the late post-stimulus period log normalized against the spectral composition of their corresponding pre-stimulus baselines. We then identified the mean log normalized early gamma and late alpha/beta band powers within each mouse and electrode location for each stimulus intensity, and then found the mean of these values across mice.

Regression of LFP power against stimulus intensity revealed significant relationships between these variables in each brain region, time and frequency range of interest, although the slope of these relationships was stronger for some than others (Figure 3). For early post-stimulus period gamma band power, A1 (figure 3C, left) exhibited the strongest response to changes in stimulus intensity with a slope of 18.037 dB/Scalar Amplitude (p = 4.099×10^−7^) and a Spearman correlation coefficient of 0.903. Thalamic Cl (Figure 3B, left) showed the next strongest response within this time and frequency span having a slope of 6.122 dB/Scalar Amplitude (p = 1.666×10^−6^) and a Spearman correlation coefficient of 0.810. The FrA (Figure 3A, left) and V1 (Figure 3D, left) both exhibited smaller slopes in early post-stimulus period, with the FrA exhibiting a slope of 3.217 dB/Scalar Amplitude (p = 8.477×10^−4^) and Spearman correlation coefficient of 0.585, while the V1 exhibited a slope of 3.559 dB/Scalar Amplitude (p = 1.649×10^−6^) and a Spearman correlation coefficient of 0.891.

For the late post-stimulus period alpha/beta band power, FrA (Figure 3A, right) showed the greatest response to stimulus intensity changes with a slope of −13.117 dB/Scalar Amplitude (p = 8.500×10^−7^) and a Spearman correlation coefficient of −0.835. The Cl (Figure 3B, right) and A1 (Figure 3C, right) exhibited weaker but still significant responses, with the Cl having a response slope of - 2.928 dB/Scalar Amplitude (p = 1.992×10^−5^) and a Spearman correlation coefficient of −0.727 and the A1 having a response slope of −4.269 dB/Scalar Amplitude (p = 1.942×10^−5^) and a Spearman correlation coefficient of −0.719. No significant relationship between stimulus volume and late post-stimulus alpha/beta band power was found in V1 (Figure 3D, right; p=0.143).

## Discussion

To assess the potential for mice as a model of the alpha/beta ERD, we recorded LFP signals in mouse frontal association cortex (FrA), thalamic intralaminar regions (Cl), as well as in primary auditory and visual cortex (A1 and V1) during passive listening and auditory go/no-go tasks with stimuli of varying intensities. This approach allowed for the isolation of electrophysiological signals created during auditory perception with excellent spatial specificity, high temporal resolution, and within the same frequency ranges employed in humans for studying alpha/beta ERDs. Our results demonstrate that mice exhibit auditory-induced decreases in alpha (8-12Hz) and beta (12-30Hz) band LFP activity at 250 to 750 ms after stimulus onset, analogous to alpha/beta ERDs previously demonstrated in humans. We also report the novel observation that the mouse Cl, a component of the intralaminar thalamus (Macchi & Bentivoglio, 1986), exhibits auditory-induced alpha/beta ERDs in a time frame comparable to those found in the mouse cortex. The ERD magnitude and relationship to auditory stimulus intensity was strongest in the association cortex area FrA, and smaller but still notable in primary auditory cortex A1 and in thalamic Cl. Finally, we found that the auditory-induced alpha/beta ERDs identified in the mouse are preceded by widespread, transient increases in broadband LFP signals, consistent with prior work showing broadband cortical signals related to local population neuronal activity (Miller, 2010; Miller et al., 2014; Mukamel et al., 2005; Mukamel et al., 2011; Nir et al., 2007). In contrast to the ERD, this early broadband signal was largest and most strongly related to auditory stimulus amplitude in A1, intermediate in Cl and FrA, and smallest in primary visual cortex V1. Taken together, these findings highlight the ability of existing methodologies to both replicate and extend prior observations in found human subjects within the more technically versatile mouse model.

### Cortical Event-Related Desynchronization

Time-frequency analysis of peristimulus LFP recordings within maximum auditory intensity trials of passive listening and go/no-go tasks revealed that mouse FrA and A1, but not V1, exhibit statistically significant decreases in alpha and beta band LFP oscillations between 250ms and 750ms post-stimulus. These results confirm that mice exhibit auditory-induced cortical desynchronizations analogous to those previously identified in humans. Auditory-induced cortical ERDs were among the first observations in humans describing the existence of the ERD electrophysiological response (Pfurtscheller & Aranibar, 1977). This original study reported decreases in cortical alpha band activity following presentation of auditory stimuli, as are reported here in mice, and further demonstrated that the ERD could be seen in the context of both passive listening and stimulus-response tasks, a finding that was also replicated here through examination of passive listening and go/no-go tasks with mice. Later work expanded on these observations, establishing that auditory-induced cortical ERDs in humans could extend into the beta band (Alegre et al., 2003; Fujioka et al., 2009; Grabska-Barwińska & Żygierewicz, 2006), a result which aligns with those in our study, and that these responses are measurable using intracranial field potential recordings (Becher et al., 2015; Christison-Lagay et al., 2025; Herman et al., 2019; Li et al., 2019) similar to those used here. Interestingly, human intracranial recordings showed that alpha/beta ERDs induced by passive auditory stimulation are significantly smaller in sleep and under anesthesia, whereas early activity in primary auditory cortex is relatively spared in these states, leading to the proposal that the alpha/beta ERD represents higher order feedback necessary for conscious perception of sensory stimuli (Hayat et al., 2022; Krom et al., 2020).

The amplitude of the mouse alpha/beta ERDs was greater in the FrA than in any other region where it was found. Similarly, regression of the mean LFP within the alpha/beta range against stimulus intensity showed the alpha/beta ERD identified in the mouse FrA was more sensitive to changes in stimulus intensity than any other region in our recordings. Interestingly, in humans auditory-induced alpha/beta ERDs are typically highest in amplitude over central or parietal, rather than frontal, brain regions (Krause et al., 1996; Krause et al., 1994; Schulter et al., 1990). Because we did not record from parietal cortex, further work will be needed to determine if the mouse ERD is larger in frontal or parietal cortex, or whether there might be differences in frontal or parietal activity reflecting differences between the neuroanatomy of rodents and humans. It is generally agreed that the rodent FrA is associated with higher brain functions including cognition and associative learning (Girard et al., 2013; Lai et al., 2018; Nakayama et al., 2015). In humans, the frontal association cortex encompasses prefrontal cortex including granular lateral prefrontal cortex (Preuss & Wise, 2022) which has been noted to be central to higher cognitive processes (Petrides, 2005). Yet it has been observed that rodents do not have a direct analog to this region (Preuss, 1995), and further it is debated to what extent rodents exhibit prefrontal cortex at all (Laubach et al., 2018). Additional research is needed to understand the cortical origin of alpha/beta ERDs in mice and their similarities to or differences from those observed in humans.

### Subcortical Event-Related Desynchronization

In this study, we show using bipolar LFP electrodes implanted in the Cl nucleus that the mouse intralaminar thalamus exhibits auditory-induced alpha/beta ERDs within the same time frame as cortical regions. This result demonstrates that subcortical regions of the brain in addition to cortical regions can exhibit alpha/beta ERDs. Further, it highlights how electrophysiological recording techniques employed in mouse models can generate new insights into the networks responsible for generating alpha/beta ERDs. Literature reporting ERDs in human subjects typically employ EEG or MEG as their primary method for examining electrophysiological activity in the brain (for review see Pfurtscheller & Lopes da Silva, 1999). These approaches are known to be effective at capturing cortical sources of activity (Kirschstein & Köhling, 2009), but the ability to ascribe subcortical sources to the signals they capture is more challenging (Krishnaswamy et al., 2017; Seeber et al., 2019). Techniques which can capture fluctuations in electrical fields deeper in the brain, such as intracranial EEG, exist and have been employed in the study of ERDs (Hayat et al., 2022). However, few studies have focused on examining ERD in subcortical regions of the human brain and none have investigated these signals in the thalamus.

What is the potential function of auditory-induced alpha/beta ERDs within the intralaminar thalamus? We discovered that the mean alpha/beta LFP power in the Cl is negatively correlated with stimulus intensity, or conversely the amplitude of the ERD increases with stimulus volume. This suggests that the networks driving alpha/beta ERDs in the intralaminar thalamus, like the cortex, is related to auditory processing. One possible role of the intralaminar thalamus in this processing is suggested by extensive previous research linking the intralaminar thalamus and Cl to arousal and acute perception through the modulation of cortical activity (Blumenfeld, 2023; Janson et al., 2021; Li et al., 2021; Schiff et al., 2013). Disruption of activity in the thalamic intralaminar region through lesions (Steriade et al., 1974) or electrical stimulation (Hunter & Jasper, 1949) have long been known to result in loss of arousal and conscious awareness. Studies have further shown that neurons in this region are sensitive to visual (Kronemer et al., 2022) or auditory stimuli (Grunwald & Eisen, 2002), and that presentation of perceived auditory stimuli but not of identical unperceived auditory stimuli is associated with increases to intralaminar thalamic neural activity (Christison-Lagay et al., 2025). Intralaminar thalamic nuclei including Cl exhibit extensive connections with the cortex (Vertes et al., 2022) and through these connections mediate synchronization and desynchronization of cortical networks (Glenn & Steriade, 1982; Gummadavelli et al., 2015; Sukov & Barth, 2001).

### Transient Broadband LFP Activity Increases

Alongside alpha/beta ERDs, we also found that presentation of auditory stimuli resulted in broadband increases to LFP activity within FrA, Cl, and A1 within 250ms of stimulus presentation. Moreover, regression of the mean gamma power in each of these regions against the intensity of the stimuli presented revealed these parameters are positively correlated, with the A1 exhibiting the greatest sensitivity to changes in stimulus intensity. These observations agree with current understanding of the mechanisms of audition and auditory cognition. The sensitivity of A1 field potentials to auditory stimulation and its necessity for audition has been well established in rodents (French, 1942; Kelly, 1980; Ross, 1965). In the FrA, neurons have been shown in single unit recordings to be sensitive to auditory stimuli (Winkowski et al., 2017), and the region as a whole has been demonstrated to mediate auditory conditioning and learning (Lai et al., 2018; Lai et al., 2012; Moczulska et al., 2013; Nakayama et al., 2015). Further, as previously discussed neurons in the Cl have been shown to respond to auditory stimulation in anaesthetized rats (Grunwerg & Krauthamer, 1992) and activity in the intralaminar thalamus mediates cortical synchrony and desynchrony (Glenn & Steriade, 1982; Gummadavelli et al., 2015) including within auditory cortical regions (Sukov & Barth, 2001). As broadband increases to LFPs have been linked to increases in activity across neuronal populations (Manning et al., 2009; Miller, 2010; Miller et al., 2014; Mukamel et al., 2005; Mukamel et al., 2011; Nir et al., 2007; Pan et al., 2011), observation of such signals within these regions, which are well-known to contribute to different aspects of audition, support the viability of LFP measures as a means of examining field potential activity in the mouse brain other than just alpha/beta ERDs.

Within the same early 0 to 250 ms time span where we found auditory-induced increases to FrA, Cl, and A1 LFP activity, we also identified increases to broadband LFP activity in V1. Regression of mean gamma band LFP power in this area of the cortex with the intensity of the stimuli presented revealed that, like in the FrA, Cl, and A1, there is also correlation between these parameters in V1. Counterintuitively, these findings are consistent with prevailing theories of visual cortex function. The visual cortex does not process visual information alone but integrates it with information from other sensory modalities (Meijer et al., 2017; Mishra et al., 2007; Vasconcelos et al., 2011). Indeed, the cross-modal sensitivity in visual cortex neurons to auditory stimuli was noted while study of the region was still nascent (Bremer, 1952; Murata et al., 1965; Spinelli et al., 1968), and more recent literature has reported auditory-induced changes to field potentials in mouse visual cortex comparable to those found here (Hattori, 2016; Land et al., 2012). The mouse V1 has been found to receive projections from the auditory cortex (Charbonneau et al., 2012; Ibrahim et al., 2016), which as previously mentioned can generate electrical field fluctuations recordable by LFP electrodes (Fernández-Ruiz et al., 2012; Makarov et al., 2010; Martín-Vázquez et al., 2013). Collectively, these literature suggest the responses to auditory stimuli found in the visual cortex in this study is not artifactual or representative of signals in other regions of the brain but reflects another example in the growing literal of cortical cross-modal processing.

### Limitations to Interpretations and Future Directions

Having established that the auditory perception-induced ERD is robust in the mouse, these findings open new avenues for research that, in future studies, could investigate the specific sources of these signals in detail, for example by monitoring or manipulating cell type-specific populations of neurons with powerful methods available in the murine models. Accurate interpretation of any electrophysiological signal in the brain demands consideration of its neuronal origin. The signals found using LFP measures of neural activity including those employed here are notoriously difficult to attribute to their specific sources. Reasons for this are numerous. Experimental validation of LFP recording methodologies confirm that the predominant source of electrical fluctuations measured by LFP electrodes are close (<250µm) to the tip (Katzner et al., 2009; Xing et al., 2009). Yet, it has been noted in some cases that volume conduction in the brain permits sources centimeters away from the electrode to also contribute (Kajikawa & Schroeder, 2011). Along these same lines terminal axonal projections have been found to generate electrical fields measurable by LFP methodologies (Fernández-Ruiz et al., 2012; Makarov et al., 2010; Martín-Vázquez et al., 2013). Such structures can originate from neuronal populations local to the electrode, but this is not necessarily the case. Further, in those cases where neuronal architecture and its electrical properties does not ambiguate LFP signal sources, modeling electrical fields can still be an issue. Single neurons are not spatially uniform sources of current flow easily representable as point sources, but instead exhibit complex patterns of electrical polarization (Martín-Vázquez et al., 2013). As a direct result of these factors, attribution of a LFP signal to its specific source requires exact understanding of the geometry of current sources in the targeted region and the location of the electrode within it (Herreras, 2016; Herreras et al., 2015). This information cannot be inferred from the methodologies performed here, but as was already mentioned, establishes a robust model for future cellular studies.

With this limitation in mind, the conclusions that can be gleaned from the results reported here are still notable. In this study we identified decreases to alpha and beta band field potential oscillations in mouse cortex following presentation of auditory stimuli characteristic of event-related desynchronization. This finding demonstrates that mice exhibit auditory-induced alpha/beta event-related desynchronizations comparable to those previously reported in humans. We further observed alpha/beta event-related desynchronizations within the mouse centrolateral thalamic nucleus following stimulus presentation. This result confirms that event-related desynchronizations can occur within subcortical regions of the brain and highlights the methodological advantages of employing mice as a model organism over humans in this area of research. Finally, we report that presentation of auditory stimuli results in widespread, broadband increases to local field potential oscillations within cortical and subcortical regions of the mouse brain that conform with modern theories of audition and auditory cognition. These data corroborate the reliability of the methodological approach used in this study. Despite the interpretive limitations of local field potential measures of neural activity, these results support the use of mice as an alternative model for investigating the role and fundamental mechanisms of event-related desynchronizations. Future studies undertaking these endeavors may find the numerous techniques available in this model organism, such as multi-unit and single-unit activity recordings or fiber photometry and optogenetics, as well as behavioral studies critical for understanding the function and purpose of event-related desynchronizations within the brain.

## Acknowledgements

This work was supported by National Institutes of Health R01 NS066974, R37 NS100901, the Mark Loughridge and Michele Williams Foundation, and the Betsy and Jonathan Blattmachr family.

